# Multi-omics phenotyping of the gut-liver axis allows health risk predictability from *in vivo* subchronic toxicity tests of a low-dose pesticide mixture

**DOI:** 10.1101/2020.08.25.266528

**Authors:** Robin Mesnage, Maxime Teixeira, Daniele Mandrioli, Laura Falcioni, Quinten Raymond Ducarmon, Romy Daniëlle Zwittink, Caroline Amiel, Jean-Michel Panoff, Emma Bourne, Emanuel Savage, Charles A Mein, Fiorella Belpoggi, Michael N Antoniou

## Abstract

Human health effects from chronic exposure to mixtures of pesticide residues are little investigated. We compared standard histopathology and serum biochemistry measures and multi-omics analyses in an *in vivo* subchronic toxicity test of a mixture of six pesticide active ingredients frequently detected in foodstuffs (azoxystrobin, boscalid, chlorpyrifos, glyphosate, imidacloprid and thiabendazole). Sprague-Dawley rats were administered with the pesticide mixture with each ingredient at its regulatory permitted acceptable daily intake. Analysis of water and feed consumption, body weight, histopathology and serum biochemistry showed little or no physiological effects from exposure to the pesticide mixture. In marked contrast, analysis of the host-gut microbiome axis using serum and caecum metabolomics revealed that nicotinamide and tryptophan metabolism were affected, which suggested the initiation of a cell danger response, including adaptation to oxidative stress. Only limited effects were detected on the caecum microbiota by shotgun metagenomics. Further analyses of *in vitro* bacterial cultures showed that growth of *Lactobacillus rhamnosus* and *Escherichia coli* strains was negatively impacted by the pesticide mixture at concentrations that were not inhibitory when exposure was to a single agent. Transcriptomics of the liver showed that 257 genes had their expression changed. Gene functions affected included those involved in the regulation of response to hormones and correlated with previously reported transcriptome changes following administration of nicotinamide. Genome-wide DNA methylation analysis of the same liver samples showed that 4255 CpG sites were differentially methylated (> 10% difference). Overall, we demonstrated that unlike standard blood biochemical and organ histological analysis, in-depth molecular profiling using a combination of high-throughput ‘-omics’ methods in laboratory animals exposed to low concentrations of pesticides reveals metabolic effects on the gut-liver axis, which can potentially be used as biomarkers for the prediction of future negative health outcomes. Our data suggest that adoption of multi-omics as part of regulatory risk assessment procedures will result in more accurate outcome measures, with positive public health implications.

## Introduction

Human exposure to pesticides has been linked to the development of a large range of diseases caused by acute intoxications or repeated occupational exposures. Occupational pesticide exposures in agricultural workers can increase cancer incidence (Kachuri et al. 2017; Leon et al. 2019) as well as the risk of developing diabetes (Juntarawijit and Juntarawijit 2018; Montgomery et al. 2008) or neurodegenerative disorders such as Parkinson’s Disease (Van Maele-Fabry et al. 2012). Populations residing in agricultural areas can also be affected. Residential proximity to pesticide use has also been associated with poorer neurodevelopment in children (Gonzalez-Alzaga et al. 2015; Gunier et al. 2017). Furthermore, studies suggest that pesticides are major contributors to the development of a wide range of chronic diseases at environmental levels of exposure in human populations (Bouchard et al. 2011; Cohn et al. 2015; Engel et al. 2011; Quirós-Alcalá et al. 2014; Raanan et al. 2016; Rauh et al. 2011; Viel et al. 2017).

The diet is a major route of exposure to pesticide residues. Dietary pesticide exposures mostly originate from the application of pesticides on to crops during cultivation (Chavarri et al. 2004), but also from the contamination of soil and water (Silva et al. 2019), as well as from post-harvest applications during storage (Jian 2019). Out of the 88,247 food samples analysed for 801 pesticides across European Union (EU) countries in 2017, 45.9% had quantifiable residues (EFSA 2019). All pesticides (169 of the 171 tested), except dimethoate and dithiocarbamates, were detected at concentrations that would result in consumption at a level lower than their regulatory permitted acceptable daily intake (ADI). It was thus concluded that current dietary pesticide exposures are unlikely to pose concerns for consumer health (EFSA 2019).

Although the evaluation of isolated chemical compounds can be a successful strategy to model toxic effects in cases of acute exposures, this approach does not realistically reflect chronic dietary exposure of human populations to mixtures of chemicals rather than single compounds (Hernandez et al. 2017). An increasing number of studies suggest that mixtures of pesticides can have toxic effects at regulatory permitted levels, which are difficult, if not impossible, to predict by testing isolated compounds (Docea et al. 2018; Lukowicz et al. 2018). Although the assessment of the cumulative exposure of pesticides on the thyroid and nervous system is currently in progress (EFSA 2019), the evaluation of effects of mixtures is not being performed in government-led programmes.

Among the different strategies to study chemical mixtures (SCHER 2012), some authors have proposed to estimate toxic effects of mixtures by simulating real-life exposures in laboratory animals (Tsatsakis et al. 2017). With this in mind, we present here a study that tested the toxic effects resulting from subchronic exposure to a mixture of the six most frequently detected pesticide residues in EU foodstuffs (EFSA 2018; EFSA 2019) in Sprague-Dawley rats. This mixture consisted of azoxystrobin (EFSA 2010), boscalid (EFSA 2014c), chlorpyrifos (EFSA 2014a), glyphosate (EFSA 2015), imidacloprid (EFSA 2008) and thiabendazole (EFSA 2014b), all combined at their respective acceptable daily intake (ADI) levels. Glyphosate is the most frequently found herbicide residue in EU foodstuffs (EFSA 2018). It is used in broad-spectrum herbicides and acts by inhibiting the plant enzyme 5-enolpyruvylshikimate-3-phosphate synthase (Schonbrunn et al. 2001). Its residues are mostly found in cereals because glyphosate-based herbicides (GBH) are frequently sprayed shortly before harvest to help crop desiccation and earlier harvest (Cessna et al. 1994). Imidacloprid and chlorpyrifos are key ingredients used in the two major categories of insecticides, neonicotinoids (Simon-Delso et al. 2015) and organophosphates (Costa 2017), respectively. The pesticide mixture also included three fungicides: namely azoxystrobin, a quinone outside inhibitor (Bartlett et al. 2002); boscalid, a succinate dehydrogenase inhibitor (Xiong et al. 2015); and thiabenbazole, a mitotic spindle distorting agent (McKellar and Scott 1990). The Stockholm Convention on Persistent Organic Pollutants (POPs), introduced in 2001 as the first global agreement to limit the use of toxic persistent chemicals with direct effects on human health, included 9 pesticides (aldrin, chlordane, DDT, dieldrin, endrin, heptachlor, hexachlorobenzene, mirex, toxaphene) among the 12 pollutants regulated in this international treaty (Porta and Zumeta 2002). This highlights the importance placed on pesticides as POPs with health implications.

The overall objective of our investigation was to identify early biochemical markers of toxicity before overt organ and system pathologies are detected. We therefore compared standard clinical and biochemical measures recommended in Organisation of Economic Co-operation and Development (OECD) guidelines for the testing of chemicals, and followed by industry and government regulatory agencies, with a multi-omics strategy. Our multi-omics approach combined shotgun metagenomics and metabolomics of caecum content, as well as serum metabolomics, liver transcriptomics, and DNA methylation profiles, to reveal the molecular mechanisms underlying metabolic changes induced by the exposure to this pesticide mixture. Our strategy takes advantage of recent developments in high-throughput “-omics technologies”, which are increasingly used to evaluate the molecular composition and dynamics of a complex system, to understand the mode of action of chemicals and to provide early biomarkers predictive of long-term health effects (Alexander-Dann et al. 2018; Taylor et al. 2018; Zampieri et al. 2018). This approach is in line with the emerging field of deep phenotyping, which seeks to measure many aspects of phenotypes and link them together to understand human health and disease.

Our findings show that unlike standard OECD blood biochemistry and organ histological analysis conducted for regulatory purposes, blood metabolomics, liver transcriptomics, and genome-wide DNA methylation analysis highlighted metabolic adaptation to oxidative stress induced by exposure to the mixture of pesticides, which could act as a predictor of long-term effects with health implications for liver function. This demonstrates the potential of in-depth molecular profiling as a measure of pesticide toxicity in animal model systems under subchronic exposure conditions. Our study is the first of its kind to directly compare standard OECD measures with a multi-omics approach. Our results suggest that the adoption of multi-omics as part of regulatory chemical risk assessment procedures will result in greater sensitivity, accuracy and predictability of outcomes from *in vivo* studies, with positive public health implications.

## Materials and methods

### Experimental animals

The experiment was conducted according to Italian law regulating the use and humane treatment of animals for scientific purposes (Decreto legislativo N. 26, 2014. Attuazione della direttiva n. 2010/63/UE in materia di protezione degli animali utilizzati a fini scientifici. – G.U. Serie Generale, n. 61 del 14 Marzo 2014). Before starting the experiment, the protocol was approved and formally authorized by the ad hoc commission of the Italian Ministry of Health (authorization N. 447/2018-PR). The experiment was conducted on female young adult Sprague-Dawley rats (8 weeks old at the start of treatment).

### Animal management

Female Sprague-Dawley rats from the Cesare Maltoni Cancer Research Center (CMCRC) breeding facility were used. Female animals were chosen in order to make the results of this investigation comparable to our previous studies (Lozano et al. 2018; Mesnage et al. 2015; Mesnage et al. 2017). The animals were generated in-house following an outbreed plan and were classified as conventional (minimal disease) status. All the experimental animals were identified by ear punch according to the Jackson Laboratory system. After weaning, and before the start of the experiment, animals were randomised in order to have at most one sister per litter of each group; homogeneous body weight within the different groups was ensured. Animals of 6 weeks of age were acclimatized for two weeks before the start of the experiment.

Rats were housed in polycarbonate cages (41×25×18 cm) with stainless wire tops and a shallow layer of white wood shavings as bedding. The animals were housed in the same room, 3 per cage, maintained at the temperature of 22±3°C and relative humidity of 50±20%. Lighting was provided by artificial sources and a 12-hour light/dark cycle was maintained. No deviations from the above-mentioned values were registered. Cages were identified by a card indicating study protocol code, experimental and pedigree numbers, and dosage group. The cages were periodically rotated on their racks to minimize effects of cage positions on animals.

### Diet and treatments

Experimental groups consisted of 12 female Sprague-Dawley rats of 8 weeks of age, treated for 90 days. Animals received ad libitum a rodent diet supplied by SAFE (Augy, France). The feed was analysed to identify possible contaminants or impurities and these were found to be below levels of detection for all substances tested (Supplementary File S1). The treatment group of animals was administered daily with a mixture of glyphosate (0.5 mg/kg bw/day), azoxystrobin (0.2 mg/kg bw/day), boscalid (0.04 mg/kg bw/day), chlorpyrifos (0.001 mg/kg bw/day), imidacloprid (0.06 mg/kg bw/day), and thiabenzadole (0.1 mg/kg bw/day), via drinking water. The concentration of pesticides in tap water to give a dose equivalent to the ADI was calculated weekly on the basis of mean body weight and mean daily water consumption. Tap water from the local mains water supplier was administered, alone or with the test compounds, to animals in glass bottles ad libitum. Every 24 hours, drinking water was discarded and the bottles cleaned and refilled. Glyphosate, azoxystrobin, boscalid, chlorpyrifos, imidacloprid, and thiabenzadole were purchased from Merck KGaA (Sigma Aldrich®, Germany) with a purity ≥ 95%.

Although the study was performed on 12 animals per group, it was originally conceived to analyse 10 animals per group with 2 animals used as a contingency in case of unexpected death as recommended by OECD guidelines for the testing of chemical toxicity. All animals survived and 12 animals per group were analysed for clinical biochemistry, histopathology, transcriptomics, methylation profiling, and shotgun metagenomics, whilst 10 animals per group were randomly chosen for the metabolomics analyses.

### Clinical observations

Animals were checked for general status three times a day, seven days a week, except non-working days when they were checked twice. Status, behaviour and clinical parameters of experimental animals were determined weekly beginning two weeks prior to commencement of treatments until the end of the experiment (at 90 days). The daily water and food consumption per cage were measured prior to the start of the experiment and weekly for the duration of the treatment. Before final sacrifice and after approximately 16 hours in a metabolic cage, water consumption was registered for each animal. Body weight of experimental animals was measured before the start of the treatment and then weekly for 90 days. All the experimental animals were weighed just before sacrifice.

### Histopathology evaluation

All sacrificed animals were subjected to complete necropsy. The gross necropsy was performed by initial physical examination of external surfaces and orifices followed by an internal *in situ* examination of tissues and organs. The examination included cranial cavity and external surfaces of the brain and spinal cord, thoracic abdominal and pelvic cavities with their associated organs and tissues, and muscular/skeletal carcass.

Samples of caecum content were collected in two vials of 100 mg each and stored at −70°C to perform evaluation of the gut microbiome. Liver and kidneys were alcohol-fixed, trimmed, processed, and embedded in paraffin wax. Sections of 3-6 μm were cut for each specimen of liver and kidneys and stained with haematoxylin and eosin. All slides were evaluated by a pathologist and all lesions of interest were reviewed by a senior pathologist.

The histopathological nomenclature of lesions adopted was in accord with international criteria; in particular non-neoplastic lesions were classified according to the international nomenclature INHAND (International Harmonization of Nomenclature and Diagnostic Criteria) and RITA (Registry of Industrial Toxicology Animal Data). Incidence of non-neoplastic lesions was evaluated with a Fisher’s exact test (one- and two-tailed; onesided results were also considered, since it is well established that only an increase in incidence can be expected from the exposure, and incidences in the control group are almost always 0).

### Blood biochemical analysis

Before sacrifice, animals were anesthetized by inhalation of a mixture of CO2/O2 (70% and 30% respectively), and approximately 7.5 ml of blood was collected from the *vena cava.* The blood collected from each animal was centrifuged in order to obtain serum, which was aliquoted into labelled cryovials and stored at −70°C. Serum biochemistry was performed under contract by IDEXX BioAnalytics (Stuttgart, Germany), an ISO 17025 accredited laboratory. Briefly, sodium and potassium levels were measured by indirect potentiometry. Albumin was measured by a photometric bromocresol green test. ALP was measured by IFCC with AMP-buffer method, glucose by Enzymatic UV-Test (hexokinase method), cholesterol by Enzymatic color test (CHOD-PAP), blood urea nitrogen by enzymatic UV-Test, gamma-glutamyl-transferase by Kinetic color test International Federation of Clinical Chemistry (IFCC), aspartate and alanine aminotransferase by kinetic UV-test (IFCC+ pyridoxal-5-phosphate), creatinine by kinetic color test (Jaffe’s method), lactate dehydrogenase by IFCC method, and triglycerides using an enzymatic color test (GPO-PAP) on a Beckman Coulter AU 480.

### DNA and RNA extraction

DNA and RNA were extracted from rat liver using the All Prep DNA/RNA/miRNA Universal Kit (Qiagen, Hilden, Germany) using the manufacturer’s instructions for ‘Simultaneous Purification of Genomic DNA and Total RNA from Animal and Human Tissues’ with no alterations. Tissue weight used was ≤ 30 mg, and samples were eluted in 30μl RNase-Free Water. RNA samples were quantified with the Nanodrop 8000 spectrophotometer V2.0 (ThermoScientific, USA) and integrity was checked using the Agilent 2100 Bioanalyser (Agilent Technologies, Waldbronn, Germany). All samples had RNA integrity numbers (RIN) ≥ 7. DNA quantity was measured using the Qubit® 2.0 Fluorometer (Life Technologies) with the dsDNA broad range reagent, followed by quality assessment using the Agilent 2200 Tapestation and Genomic DNA screentape (Agilent Technologies). Samples displayed high molecular weight with DINs (DNA integrity score) ranging from 7.5 to 10, and average concentrations from 370 ng/μL. All samples showed a majority of high molecular weight material and all were taken forward for processing.

### Transcriptomics analysis

For RNA-seq transcriptomics, 100ng of total RNA was used for each sample for library preparation. mRNA libraries were prepared using the NEBNext® Poly(A) mRNA Magnetic Isolation Module in combination with the NEBNext® Ultra™ II Directional RNA Library Prep Kit and indexed with NEBNext® Multiplex Oligos for Illumina® (96 Index Primers) (New England Biolabs, Ipswich, Massachusetts, USA). Fragmentation was carried out using incubation conditions recommended by the manufacturer for samples with a RIN ≥7 to produce RNA sizes ≥ 300bp (94°C for 10 minutes) with first strand synthesis carried out by incubation at 42°C for 50 minutes. Modification of the manufacturer’s conditions was used when enriching Adaptor Ligated DNA for libraries with a size of 300bp and 13 cycles of PCR were performed for final library enrichment. Resulting libraries were quantified using the Qubit 2.0 spectrophotometer (Life Technologies, California, USA) and average fragment size assessed using the Agilent 2200 Tapestation (Agilent Technologies, Waldbronn, Germany). Sample libraries were combined in equimolar amounts into a single pool. The final library pool was sequenced twice on the NextSeq500 at 1.1pM and 75bp paired-end reads were generated for each library using the Illumina NextSeq®500 v2.5 High-output 150 cycle kit (Illumina Inc., Cambridge, UK). A total of 319,920,579 reads (average of 13,330,024 ± 3,068,802 reads per sample) were generated for the 24 liver samples. The raw data from the transcriptomics analysis is available at the SRA accession number [to be provided].

### Reduced representation bisulfite sequencing

A total of 100ng of total DNA was diluted and processed using the Premium Reduced Representation Bisulfite Sequencing (RRBS) Kit (Diagenode, Denville, NJ, USA) as per the manufacturer’s instructions. Briefly, DNA was digested with *MspI* prior to end repair, adapter ligation and size selection. Products were then amplified by qPCR and pooled in equal amounts. Pooled libraries were bisulphite converted and PCR enriched following a second qPCR amplification. Libraries were quantified using the Qubit® 2.0 Fluorometer (Life Technologies) followed by quality assessment using the Agilent 2200 Tapestation and DS1000 screentape (Agilent Technologies). Pooled libraries were loaded at 1.1M with 20% standard PhiX library (Illumina, CA, USA) and sequenced to 75 base pair single end on a NextSeq 500 (Illumina, CA, USA). Data was aligned to the rat reference genome Rn6 with Bismark (Krueger and Andrews 2011). A total of 407,904,185 reads (average of 16,996,008 ± 4,648,420 reads per sample) were generated for the 24 liver samples. The raw data from the RRBS analysis is available at the SRA accession number [to be determined].

### Metabolomics

Metabolomics analysis was conducted under contract with Metabolon Inc. (Durham, NC, USA). Samples were prepared using the automated MicroLab STAR® system from Hamilton Company. Proteins were precipitated with methanol under vigorous shaking for 2 min (Glen Mills GenoGrinder 2000), followed by centrifugation. Samples were placed briefly on a TurboVap® (Zymark, USA) to remove the organic solvent. The sample extracts were stored overnight under nitrogen before preparation for analysis. The resulting extract was analysed on four independent instrument platforms: two different separate reverse phase ultrahigh performance liquid chromatography-tandem mass spectroscopy analysis (RP/UPLC-MS/MS) with positive ion mode electrospray ionization (ESI), a RP/UPLC-MS/MS with negative ion mode ESI, as well as by hydrophilic-interaction chromatography (HILIC)/UPLC-MS/MS with negative ion mode ESI.

All UPLC-MS/MS methods utilised a Waters ACQUITY ultra-performance liquid chromatography (UPLC) and a Thermo Scientific Q-Exactive high resolution/accurate mass spectrometer interfaced with a heated electrospray ionization (HESI-II) source and Orbitrap mass analyser operated at 35,000 mass resolution. The sample extract was dried and then reconstituted in solvents compatible to each of four methods used. Each reconstitution solvent contained a series of standards at fixed concentrations to ensure injection and chromatographic consistency. One aliquot was analysed using acidic positive ion conditions, chromatographically optimised for more hydrophilic compounds. In this method, the extract was gradient eluted from a C18 column (Waters UPLC BEH C18-2.1×100 mm, 1.7 μm) using water and methanol, containing 0.05% perfluoropentanoic acid (PFPA) and 0.1% formic acid (FA). Another aliquot was also analysed using acidic positive ion conditions, chromatographically optimised for more hydrophobic compounds. In this method, the extract was gradient eluted from the same afore mentioned C18 column using methanol, acetonitrile, water, 0.05% PFPA and 0.01% FA and was operated at an overall higher organic content. Another aliquot was analysed using basic negative ion optimised conditions using a separate dedicated C18 column. The basic extracts were gradient eluted from the column using methanol and water, with 6.5mM ammonium bicarbonate at pH 8. The fourth aliquot was analysed via negative ionization following elution from a HILIC column (Waters UPLC BEH Amide 2.1×150 mm, 1.7 μm) using a gradient consisting of water and acetonitrile with 10mM ammonium formate, pH 10.8. The MS analysis alternated between MS and data-dependent MSn scans using dynamic exclusion. The scan range varied slightly between methods but covered 70-1000 m/z.

Raw data was extracted, peak-identified and QC processed using Metabolon’s hardware and proprietary software. Compounds were identified by comparison to library entries of purified standards or recurrent unknown entities. Biochemical identifications are based on three criteria: retention index within a narrow retention time/index (RI) window of the proposed identification, accurate mass match to the library +/− 10 ppm, and the MS/MS forward and reverse scores between the experimental data and authentic standards. Peaks were quantified using area-under-the-curve. Raw data is available in Metabolights with accession number MTBLS138.

### Shotgun metagenomics

DNA was extracted from 100 mg caecum content using the Quick-DNA Fecal/Soil Microbe Miniprep Kit (Zymo Research, Irvine, CA, USA) with minor adaptations from manufacturer’s instructions (Ducarmon et al. 2020). Adaptations were: 1. bead beating was performed at 5.5 m/s for three times 60 seconds using a Precellys 24 homogeniser (Bertin Instruments, Montigny-le-Bretonneux, France) and 2.50 μl elution buffer was used to elute the DNA, following which the eluate was run over the column once more to increase DNA yield. One negative extraction control (no sample added) and one positive extraction control (ZymoBIOMICS Microbial Community Standard; Zymo Research) were processed in parallel during the DNA extraction procedures and subsequently sequenced. DNA was quantified using the Qubit HS dsDNA Assay kit on a Qubit 4 fluorometer (Thermo Fisher Scientific, Horsham, UK).

Shotgun metagenomics was performed by GenomeScan (Leiden, The Netherlands). The NEBNext® Ultra II FS DNA module (cat# NEB #E7810S/L) and the NEBNext® Ultra II Ligation module (cat# NEB #E7595S/L) were used to process the samples. Fragmentation, A-tailing, and ligation of sequencing adapters of the resulting product was performed according to the procedure described in the NEBNext Ultra II FS DNA module and NEBNext Ultra II Ligation module Instruction Manual. The quality and yield after sample preparation were measured with the Fragment Analyzer (Table 2 and Appendices). The size of the resulting product was consistent with the expected size of approximately 500-700 bp. Clustering and DNA sequencing using the NovaSeq6000 (Illumina inc.) was performed according to manufacturer’s protocols. A concentration of 1.1 nM of DNA was used. NovaSeq control software NCS v1.6 was used.

The shotgun metagenomics data was pre-processed using the pre-processing package v0.2.2 for cleaning metagenomes sequenced on the Illumina HiSeq (https://anaconda.org/fasnicar/preprocessing). In brief, this package (1) concatenates all forward reads into one file and all reverse reads into another file, (2) uses trim_galore to remove Illumina adapters, trim low-quality positions and unknown position (N) and discard low-quality (quality <20 or >2 Ns) or too short reads (< 75bp), (3) removes contaminants (phiX and rat genome sequences), (4) sorts and splits the reads into R1, R2, and UN sets of reads. Raw data is available in the SRA with accession number PRJNA609596.

Cleaned shotgun metagenomics reads were then processed for taxonomic and pathway profiling. Since there is no gold standard for computational analyses of shotgun metagenomics, we used a combination of approaches. In order to use as many DNA reads as possible, we inferred the taxonomy with the RefSeq database on the metagenomics RAST server (Keegan et al. 2016). We also inferred the taxonomy with a variety of tools. We used IGGsearch (iggdb_v1.0.0_gut database), which used taxa specific marker genes and circumvents problems associated with the amplification of single amplicons such as in the case of the 16S rRNA gene (Nayfach et al. 2019). Taxonomic composition was also derived with MetaPhlAn version 2.9 (Truong et al. 2015) and Kaiju 1.0.1 (Menzel et al. 2016).

### *In vitro* study of bacterial growth

Azoxystrobin, boscalid, chlorpyrifos, imidacloprid and thiabendazole were diluted in dimethylsulfoxide (DMSO) (Merck, Feltham, UK) to obtain stock solutions. Glyphosate was diluted in water. Bacterial strains were provided by the Université de Caen Microbiologie Alimentaire (Caen, France) culture collections (Table S1). The broth dilution method was used to determine how pesticides modify the bacterial growth under aerobic conditions. Bacteria from overnight broth cultures were re-suspended to OD_600nm_ = 0.3 (*Lactobacillus rhamnosus*) or OD_600nm_ = 0.2 (*Escherichia coli*) and further diluted 1,000-fold in MRS without peptone for *L. rhamnosus,* or ABTG for *E. coli* broth, to obtain approximately 10^5^ CFU/ml as confirmed in each experiment by plating cell suspension on agar plates incubated aerobically at 37 °C. Plates were incubated under aerobic conditions at 37°C and inspected after 24h and 48h. All experiments were performed in triplicate.

### Statistical analysis

The metabolome data analysis was performed using R version 3.9 (Team 2019). Peak area values were median scaled, log transformed, and any missing values imputed with sample set minimums, both on a per biochemical basis, and separately for each metabolome dataset. Statistical significance was determined using a Welch’s two-sample t-test adjusted for multiple comparisons with FDR methods using the R package ‘qvalue’ version 2.17.0 (Storey and Tibshirani 2003). For the shotgun metagenomics, a compositional data analysis approach was used since gut metagenomics datasets are typically zero-inflated (Knight et al. 2012). We used ALDEx version 2 (ALDEx2) for differential (relative) abundance analysis of proportional data (Fernandes et al. 2013). Statistical analysis for taxa abundance was performed on a dataset corrected for asymmetry (uneven sequencing depths) using the *inter-quartile log-ratio* method, which identifies features with reproducible variance. Given the relatively small number of samples in this study, we assessed statistical significance using a Wilcoxon test, with p-values adjusted for multiple comparisons with the FDR approach. A multivariate analysis consisting in a non-metric multidimensional scaling (NMDS) plot of Bray-Curtis distances between samples. Statistical significance of the sample clustering was evaluated with a permutational ANOVA (PERMANOVA) analysis on the Bray-Curtis distances with *adonis(*) from vegan v2.4-2.

We also used orthogonal partial least squares discriminant analysis (OPLS-DA) to evaluate the predictive ability of each omics approach. OPLS-DA is an extension of PLS methods, which includes an orthogonal component distinguishing the variability corresponding to the experimental perturbation (here the effects of the pesticide mixture) from the portion of the data that is orthogonal; that is, independent from the experimental perturbation. The R package ropls version 1.20.0 was used (Thévenot et al. 2015). This algorithm uses the nonlinear iterative partial least squares algorithm (NIPALS). Prior to analysis, experimental variables were centred and unit-variance scaled. Since PLS-DA methods are prone to overfitting, we assessed the significance of our classification using permutation tests (permuted 1,000 times).

RNA-seq data was analysed with Salmon (Patro et al. 2017). This tool was used to quantify transcript abundance by mapping the reads against a reference transcriptome (Ensembl Release Rattus Norvegicus 6.0 cDNA fasta). Mapping rate was 82.0 ± 4.4% on a rat transcriptome index containing 31,196 targets. The Salmon output was then imported in R version 3.9. (Team 2019) using the Bioconductor package tximport. We created a transcript database containing transcript counts, which was used to perform a differential gene expression analysis using DESeq2 (Love et al. 2014). We finally used goseq to perform a gene ontology analysis accounting for transcript length biases (Young et al. 2010). We also compared our transcriptome findings to a list of gene expression signatures collected from various rat tissues after treatments with various drugs using the drugMatrix toxicogenomics database (Ganter et al. 2006) with EnrichR (Kuleshov et al. 2016).

DNA methylation calls from RRBS data were extracted with Bismark (Krueger and Andrews 2011). The output from Bismark was then imported in R and analysed with Methylkit (Akalin et al. 2012). DNA methylation calls were annotated using RefSeq gene predictions for rats (rn6 release) with the package genomation (Akalin et al. 2014). Other annotations were retrieved using the genome wide annotation for rat tool org.Rn.eg.dbR package version 3.8.2. Statistical analysis was performed with logistic regression models fitted per CpG using Methylkit functions. P-values were adjusted to Q-values using SLIM method (Wang et al. 2011).

Statistical analyses of *in vitro* tests on bacterial growth were performed using GraphPad Prism version 8.0.1 (GraphPad Software, Inc, CA, USA). Differences between treatment groups at different concentrations and the negative control were investigated using Kruskal-Wallis one-way ANOVA with Dunn’s multiple comparison post-test.

## Results

The aim of this study was to test the toxicity *in vivo* of a mixture of pesticides, the residues of which are amongst those most frequently found in the EU food chain. A group of 12 twelve female Sprague-Dawley rats were administered with a combination of azoxystrobin, boscalid, chlorpyrifos, glyphosate, imidacloprid and thiabendazole for 90-days via drinking water to provide the EU ADI and compared to an equivalent control group receiving plain drinking water (Figure 1A and 1B). No significant differences were observed between the treatment and control groups of animals in terms of water consumption (Figure 1C), feed consumption (Figure 1D) and mean body weight (Figure 1E) during the 90-day treatment period. Histological analysis showed there was a nonsignificant increase in the incidence of liver and kidney lesions (Figure 2A and 2B). A serum biochemistry analysis only detected a small reduction in creatinine levels (Figure 2C).

**Figure 1.**
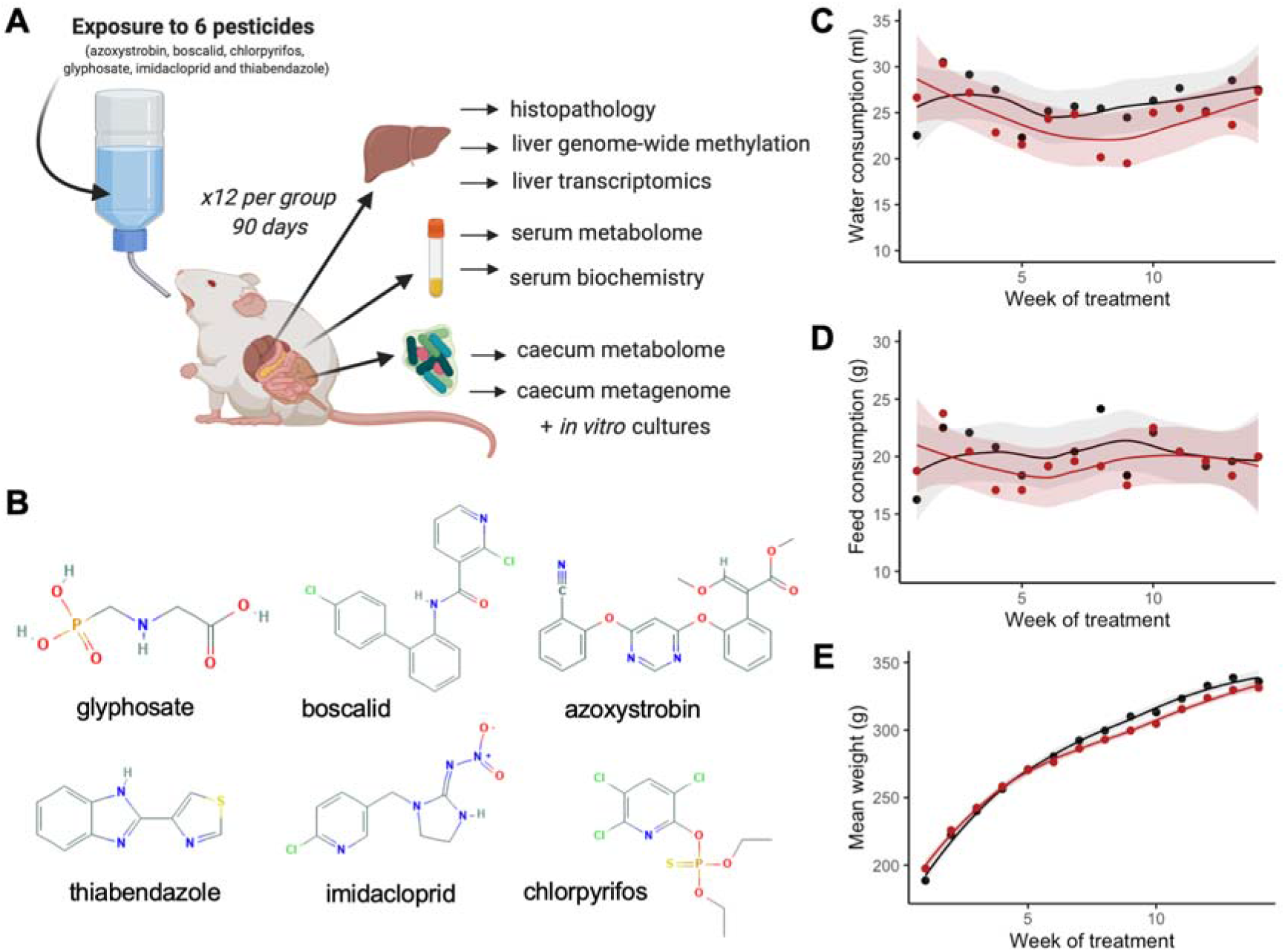
General toxicity assessment of a mixture of glyphosate, azoxystrobin, boscalid, chlorpyrifos, imidacloprid, and thiabenzadole at their acceptable daily intake in Sprague-Dawley rats. **(A)** Study design. Groups of 12 female rats were administered via drinking water with a mixture of 6 pesticides at the EU ADI for 90-days. Analyses following sacrifice are also shown. (Illustration created with BioRender.com) **(B)** Molecular structures of pesticide active ingredients tested. (Chemical structure from pubchem.com). Water consumption **(C)**, feed consumption **(D)** and body weights with their 95% confidence interval band **(E)** remained unchanged (controls, black; pesticide-exposed, red).

**Figure 2.**
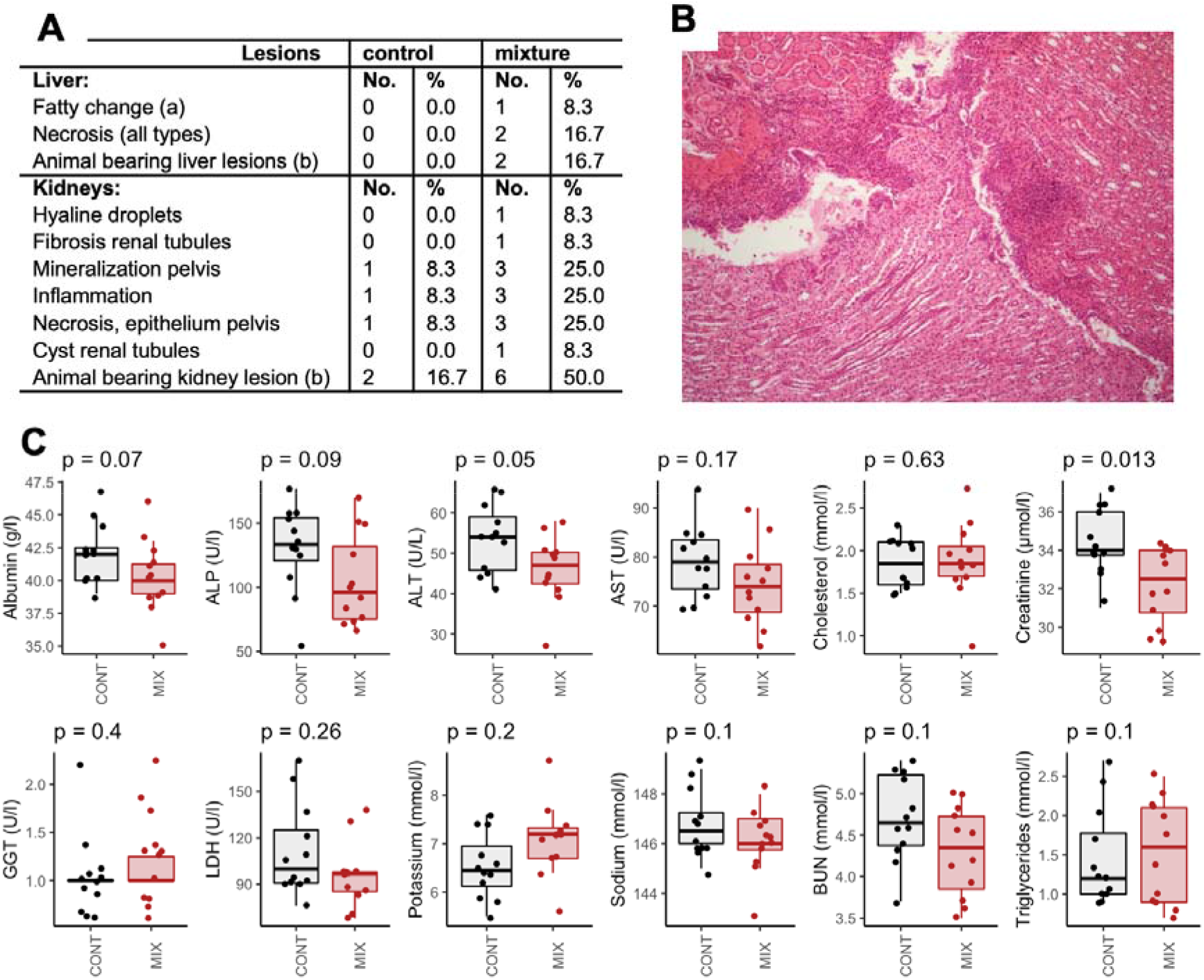
Analysis of regular clinical and biochemical markers provided limited insight into the effects of a mixture of pesticides at their acceptable daily intake in Sprague-Dawley rats. Incidence in signs of anatomical pathologies in liver and kidneys **(A).** Focal inflammation of moderate to severe grade localised in the pelvic area in a rat exposed to the pesticide mixture; magnification 100X **(B)**. Standard serum biochemistry analysis showed only a decrease in creatinine levels **(C)**.

In order to obtain insight into the possible chronic effects of this pesticide mixture, we used high-throughput molecular profiling techniques to search for changes, which could predict the development of disease. We built OPLS-DA models to assess the predictive ability of the different omics approaches used in this study. Serum metabolomics (pR2Y = 0.003, pQ2 = 0.001), liver transcriptomics (pR2Y = 0.10, pQ2 = 0.002), and to a lesser extent the caecum metabolomics (pR2Y = 0.16, pQ2 = 0.003), discriminated the pesticide-treated group from the concurrent control group (Table 1). Shotgun metagenomics in caecum and genome-wide methylation in liver did not significantly discriminate between the two experimental groups.

**Table 1.** Predictive ability of high-throughput omics approaches to evaluate the effects of a pesticide mixture in rats. OPLS-DA models were performed for each set of omics data. We show estimates of the total explained variation (R2X), variations between the different groups (R2Y) and the average prediction capability (Q2). We assessed the significance of our classification using permutation tests. The number of variables investigated (n) is also shown. New estimates of R2Y and Q2 values were calculated from this 1,000 times permuted dataset (p-values pR2Y and pQ2 for permuted R2Y and Q2, respectively).

| Omics technology | n | R2X | R2Y | Q2 | pR2Y | pQ2 |
| --- | --- | --- | --- | --- | --- | --- |
| serum metabolome | 749 | 0.34 | 0.99 | 0.80 | 0.003 | 0.001 |
| caecum metabolome | 744 | 0.33 | 0.87 | 0.49 | 0.16 | 0.003 |
| caecum metagenome | 2466 | 0.72 | 0.99 | 0.71 | 0.85 | 0.3 |
| liver transcriptome | 18170 | 0.48 | 0.97 | 0.49 | 0.10 | 0.002 |
| liver genome-wide methylation | 136555 | 0.06 | 0.88 | 0.00 | 0.4 | 1 |

Analysis of the host-gut microbiome axis using metabolomics revealed effects on the tryptophan–nicotinamide pathway (Table 2). A decrease in serum tryptophan levels, and its breakdown product indoleacetate, suggested that this amino acid is channeled to nicotinamide synthesis. The three metabolites nicotinamide N-oxide, 1-methylnicotinamide and nicotinamide all increased. In addition, the decrease in pyridoxal can also be linked to changes in nicotinamide metabolism (Table 2), since it is an important co-factor necessary for the synthesis of nicotinamide adenine dinucleotide.

**Table 2.**
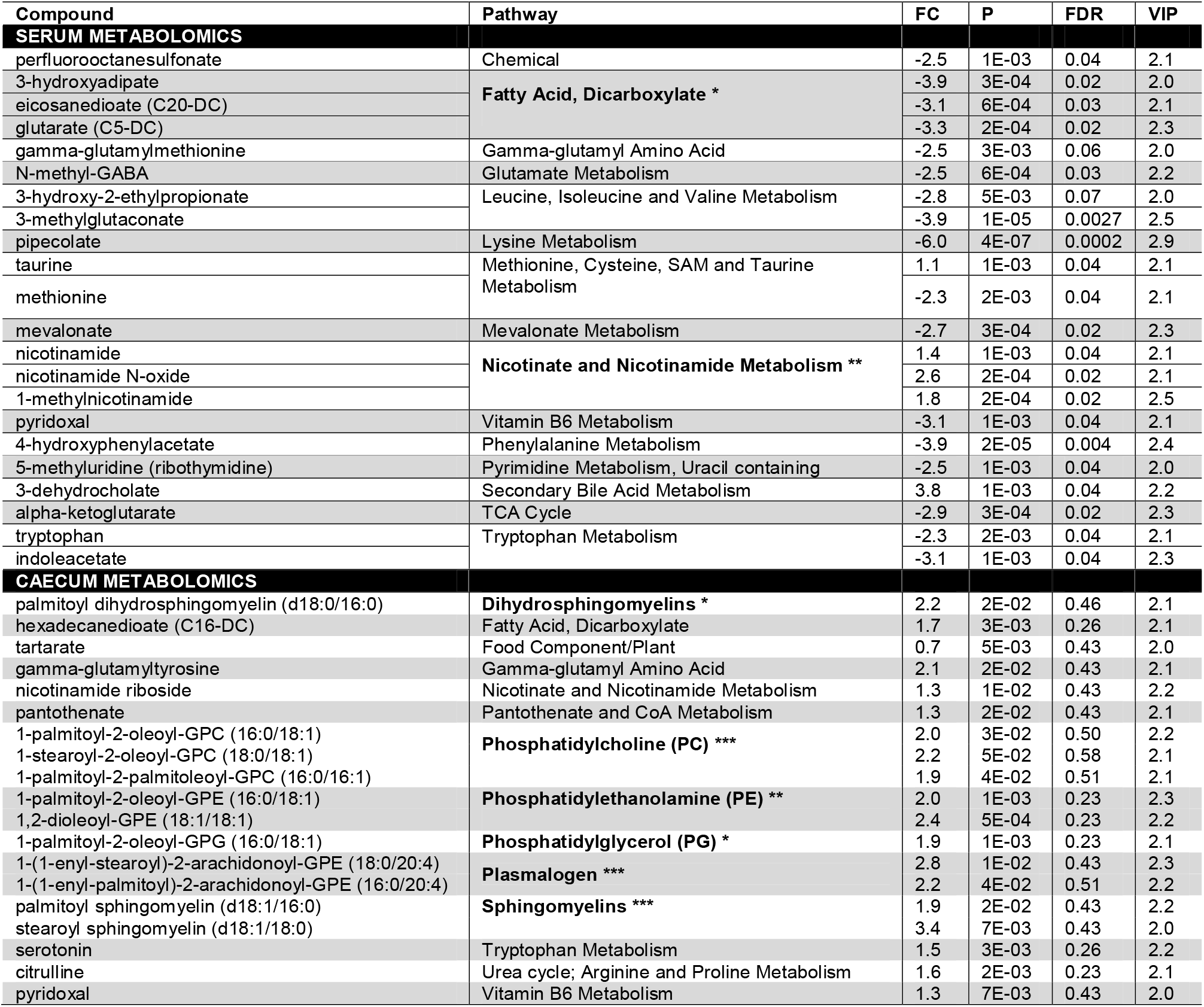
Serum and caecum metabolomics of host-gut microbiota interactions in rats exposed to a pesticide mixture reveals alterations in multiple metabolic pathways. We presented fold changes (FC) for the metabolites which were found to have their variable importance in projection (VIP) scores > 2 in the OPLS-DA analyses. P-values from a Welch’s t-test (P) are presented with the FDR. The statistical significance of a pathway enrichment analysis (bold character, p < 0.05) is also presented (* p < 0.05 ; ** p < 0.01 ; *** p < 0.001).

**Table 3.** Differentially methylated CpG sites located at gene promoters. Reduced Representation Bisulfite Sequencing was performed on liver samples. Differentially methylated CpG sites present in promoters were filtered and their variations summarized.

| chr | coordinates | strand | p | FDR | % diff | gene.name |
| --- | --- | --- | --- | --- | --- | --- |
| chr7 | 2787189 | - | 5E-57 | 5E-53 | 15.4 | coenzyme Q10A |
| chr3 | 57717495 | + | 8E-55 | 5E-51 | -11.6 | cytochrome b reductase 1 |
| chr14 | 4250209 | + | 8E-50 | 1E-46 | 13.5 | uncharacterized LOC102551276 |
| chr7 | 138705521 | + | 4E-39 | 2E-36 | 11.1 | PC-esterase domain containing 1B |
| chr1 | 219852329 | - | 6E-33 | 2E-30 | 10.8 | leucine rich repeat and fibronectin type III domain containing 4 |
| chrX | 134538342 | + | 3E-32 | 7E-30 | -11.4 | similar to CXXC finger 5 |
| chr5 | 159427478 | + | 2E-28 | 3E-26 | -10.1 | peptidyl arginine deiminase 2 |
| chr15 | 33120272 | + | 4E-27 | 7E-25 | -11.6 | RRAD and GEM like GTPase 2 |
| chr3 | 147818650 | - | 3E-25 | 5E-23 | -10.3 | tribbles pseudokinase 3 |
| chr17 | 80793952 | + | 1E-24 | 2E-22 | 19.2 | uncharacterized LOC102550536 |
| chr20 | 3350268 | + | 3E-23 | 4E-21 | 16.9 | alpha tubulin acetyltransferase 1 |
| chr19 | 40616367 | + | 2E-21 | 2E-19 | -11.7 | uncharacterized LOC103694327 |
| chr20 | 4362151 | + | 1E-19 | 1E-17 | 12 | advanced glycosylation end product-specific receptor |
| chr13 | 53571396 | - | 4E-19 | 4E-17 | 13.6 | uncharacterized LOC102547588 |
| chr1 | 87044466 | + | 5E-17 | 3E-15 | 16.8 | galectin 7 |
| chr8 | 70015572 | - | 1E-16 | 9E-15 | 13.3 | uncharacterized LOC102548470 |
| chr2 | 53310072 | - | 5E-16 | 3E-14 | 10.8 | growth hormone receptor |
| chr6 | 28234694 | + | 4E-15 | 2E-13 | 11 | DNA methyltransferase 3 alpha |
| chr5 | 154524190 | + | 1E-11 | 4E-10 | -11.7 | E2F transcription factor 2 |
| chr14 | 45398591 | - | 1E-11 | 3E-10 | -11.1 | uncharacterized LOC102552762 |
| chr10 | 107157939 | + | 2E-11 | 6E-10 | -10.7 | uncharacterized LOC103693482 |
| chr1 | 220883570 | - | 2E-09 | 3E-08 | -13.3 | melanoma-associated antigen G1-like |
| chr5 | 172527225 | - | 9E-08 | 1E-06 | 10.3 | uncharacterized LOC108351067 |
| chr9 | 4547982 | + | 9E-06 | 6E-05 | 10.7 | uncharacterized LOC108351871 |

Since the gut microbiome has known roles on nicotinamide metabolism, we analysed the composition and function of the caecum microbiome through shotgun metagenomics and metabolomics. The most discriminant metabolites between control and treatment groups of animals are glycerophospholipids, which accumulated in the caecum (Table 2). In addition, the increased levels of serotonin, which has both neurotransmitter and hormone functions and is synthesized from tryptophan, as well as the increased levels of pyridoxal and nicotinamide riboside, suggested that the effects of the pesticide mixture in the caecum microbiome can be linked to the observed disruptions in serum metabolite levels (Table 2).

Shotgun metagenomics (Figure 3A) showed that the caecum metagenome of the Sprague-Dawley rats was dominated by Firmicutes and Bacteroidetes. No species were detected in a negative extraction control, which was included to ensure that no bacterial contamination was introduced by laboratory reagents and procedures. PERMANOVA analysis of the differences between Bray-Curtis distances calculated from the abundance determined with IGGsearch did not show a significant effect of the pesticide treatment (Figure 3C). No differences in species abundance were identified with a range of shotgun metagenomics taxonomic classifiers such as IGGsearch (Excel Table S3), MetaPhlan2 (Excel Table S4) or Kaiju (Excel Table S5). There was a large intragroup variability by comparison to the variability observed between the two groups of rats (species-level operational taxonomic units of IGGsearch shown in Figure 3B), which prevented reaching definitive conclusions. This was also the case when we measured pathway abundances for tryptophan (Figure 3D) and nicotinamide metabolism (Figure 3E).

**Figure 3.**
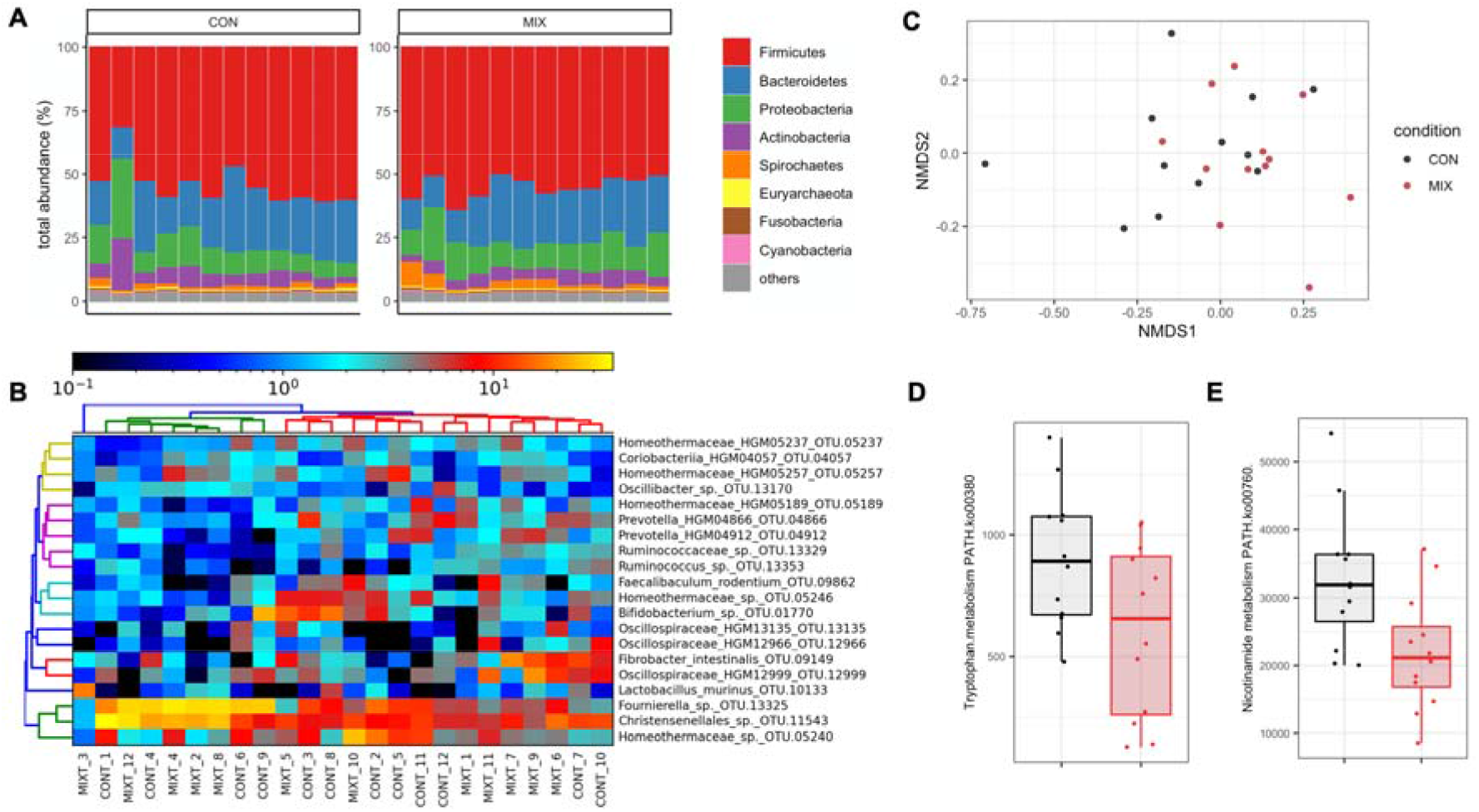
Shotgun metagenomics shows no alterations in the caecum microbiota composition upon exposure to the mixture of 6 pesticides. **(A)** Gut caecum microbiota composition profiles at the phylum level. **(B)** Classification of samples from the most frequently found species-level operational taxonomic units with IGGsearch failed to show any alterations in different bacterial populations in response to the pesticide mixture. **(C)** Principal coordinates analysis plot using the NMDS ordination of Bray-Curtis distances. Pathway analysis shows reduction in tryptophan **(D)** and nicotinamide **(E)** metabolism potential.

We then assessed if the mixture of pesticides could have effects on two strains of *Escherichia coli* and two strains of *Lactobacillus rhamnosus,* which are found in the human intestinal microbiota (Figure 4). When the six pesticide active ingredients were tested individually, they did not affect the growth of the four bacterial strains (Figure S1 and S2). However, when the bacterial strains were exposed to the mixture of all six pesticides, growth inhibition was detected in a strain-dependent manner. Although *L. rhamnosus* LB5 (Figure 4A) and *E. coli* EC4 (Figure 4C) had their growth inhibited by the pesticide mixture, *L. rhamnosus* LB6 (Figure 4B) and *E. coli* EC2 (Figure 4D) were not affected.

**Figure 4.**
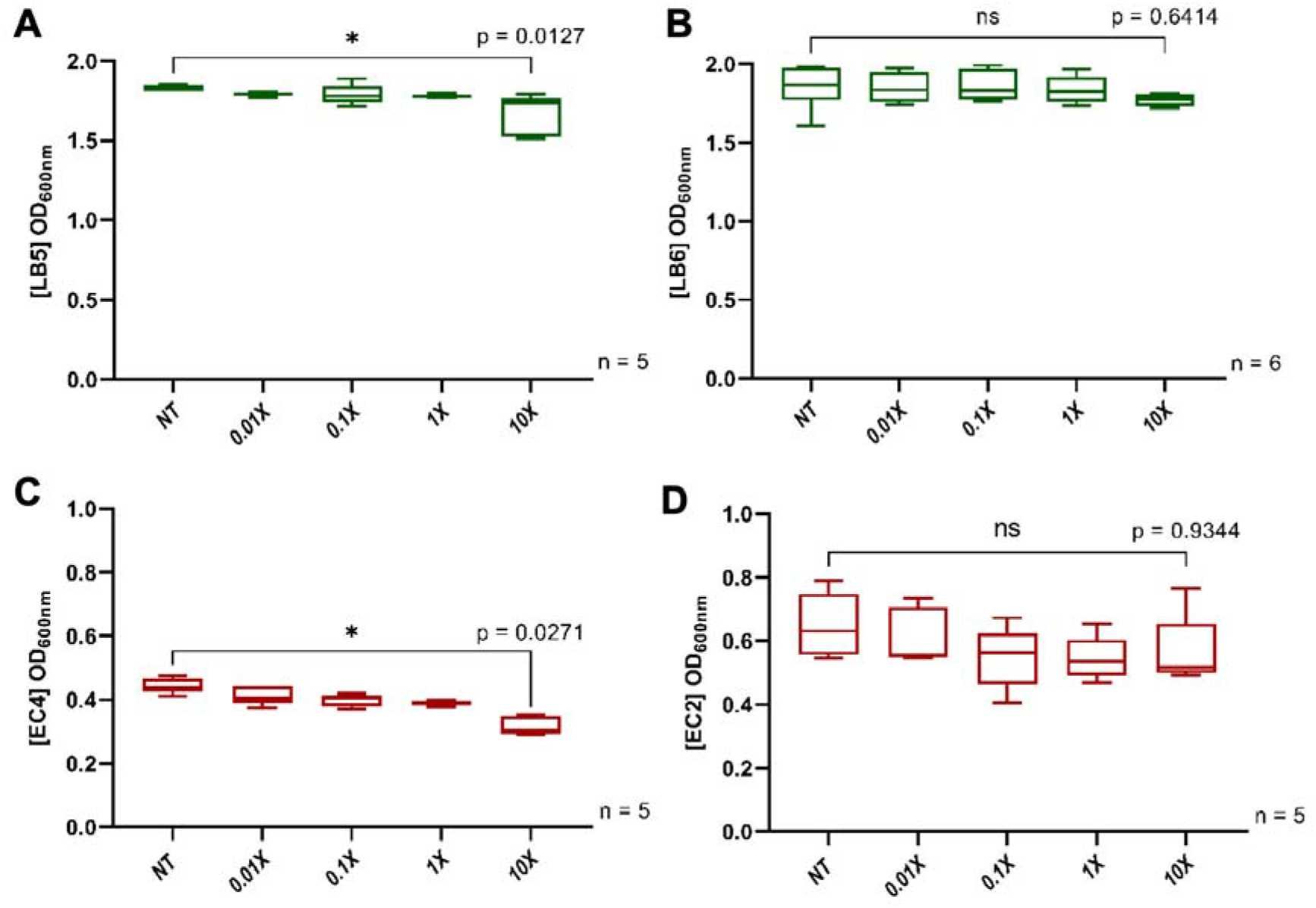
Effects of the pesticide mixture on *Lactobacillus rhamnosus* and *Escherichia coli* in vitro. Bacterial growth of the strain *Lactobacillus rhamnosus* (LB5) **(A)** is significantly inhibited after an exposure at 10 times the ADI (10 X) for the mixture while another strain of *L. rhamnosus* (LB6) **(B)** was not inhibited at the same dose. Similarly, *Escherichia coli* (EC4) growth **(C)** was inhibited at lower doses in comparison with the other *E. coli* strain (EC2) **(D)**.

Mammals mostly produce the vitamin nicotinamide (a form of vitamin B3) from tryptophan in the liver, before it is distributed to non-hepatic tissues. In order to understand if the changes in the gut and blood metabolome we observed are associated with disturbance in liver function, we compared transcriptome profiles in the liver of the two groups of rats by RNA sequencing (Figure 5). The results showed that the expression of 121 genes was increased and 117 genes had their expression decreased (Excel Table S6) by exposure to the pesticide mixture (adj-p < 0.05). A total of 96 Gene Ontology (GO) terms were enriched among our differentially expressed genes. Most of them were involved in the regulation of response to hormones (adj-p = 0.0003). Interestingly, the two genes with the lowest p-values had their expression increased and coded the Nuclear Receptor Subfamily 1 Group D Member 1 (NR1D1, adj-p = 3.2e-17) and 2 (NR1D2, adj-p = 1.9e-18), which play a role in carbohydrate and lipid metabolism, as well as in in regulation of circadian rhythms. We finally compared our transcriptome findings to a list of gene expression signatures collected from various rat tissues after treatments with various drugs from the drugMatrix toxicogenomics database (Table S2). The alterations in the liver transcriptome we observed correlated well with those of livers from a study in which nicotinamide was administered to rats at a dose of 750 mg/kg (adj-p = 1.6e-7). This correlates well with the changes in the biochemical composition of the serum-gut microbiome axis.

**Figure 5.**
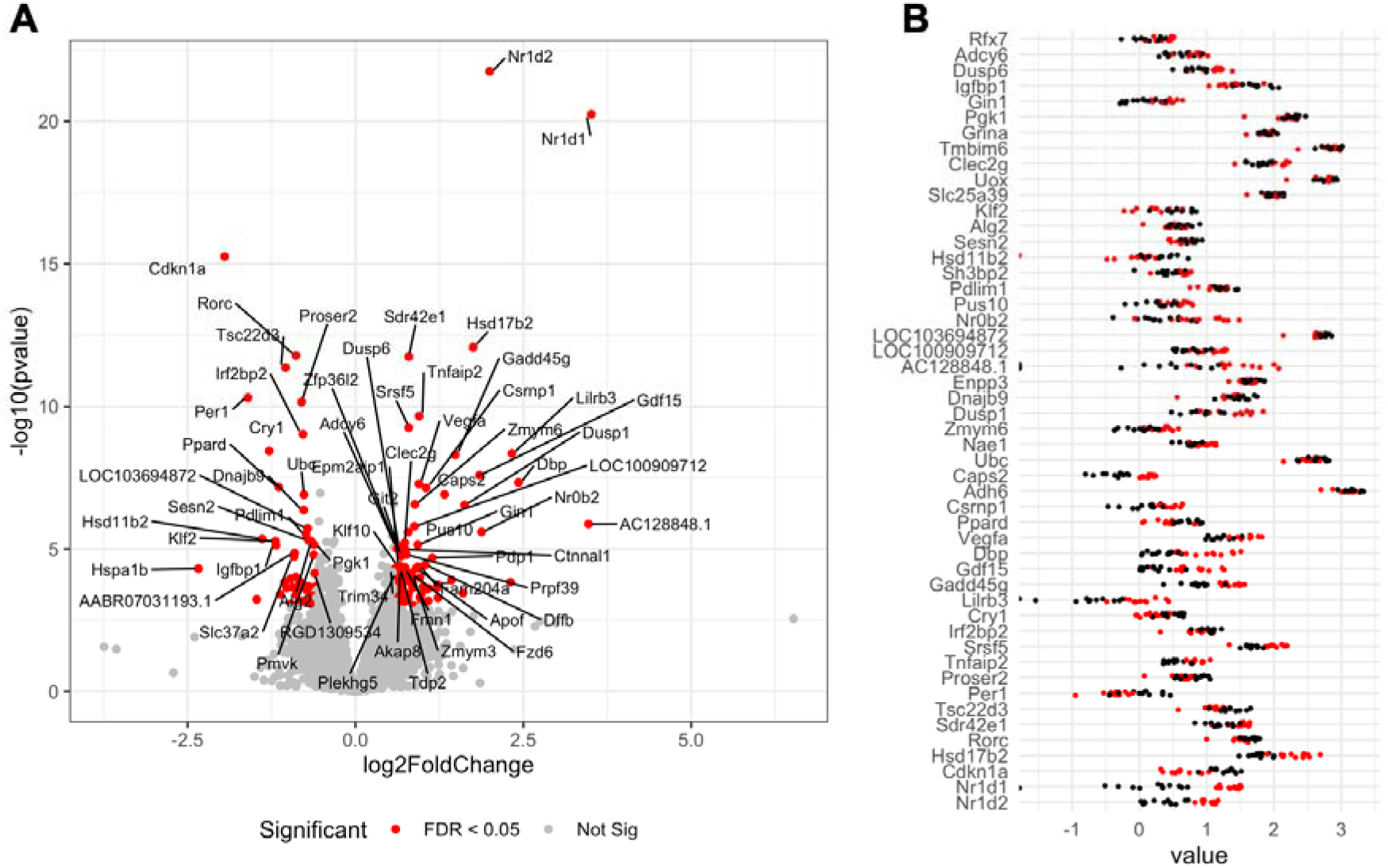
Liver transcriptome of Sprague-Dawley rats exposed for 90 days to the mixture of 6 pesticides. Genes were considered as differentially expressed if their count were found to be statistically significant after an analysis with DESeq2. **A**. A volcano plot showing the fold changes and statistical significance in the expression of genes affected by exposure to the pesticide mixture. **B.** The effect size for the 50 most affected transcripts. Log10 normalised abundances from the DESeq2 analysis were used to facilitate the visualization of differences (red dots, pesticide treated; black dots, untreated controls).

**Figure 6.**
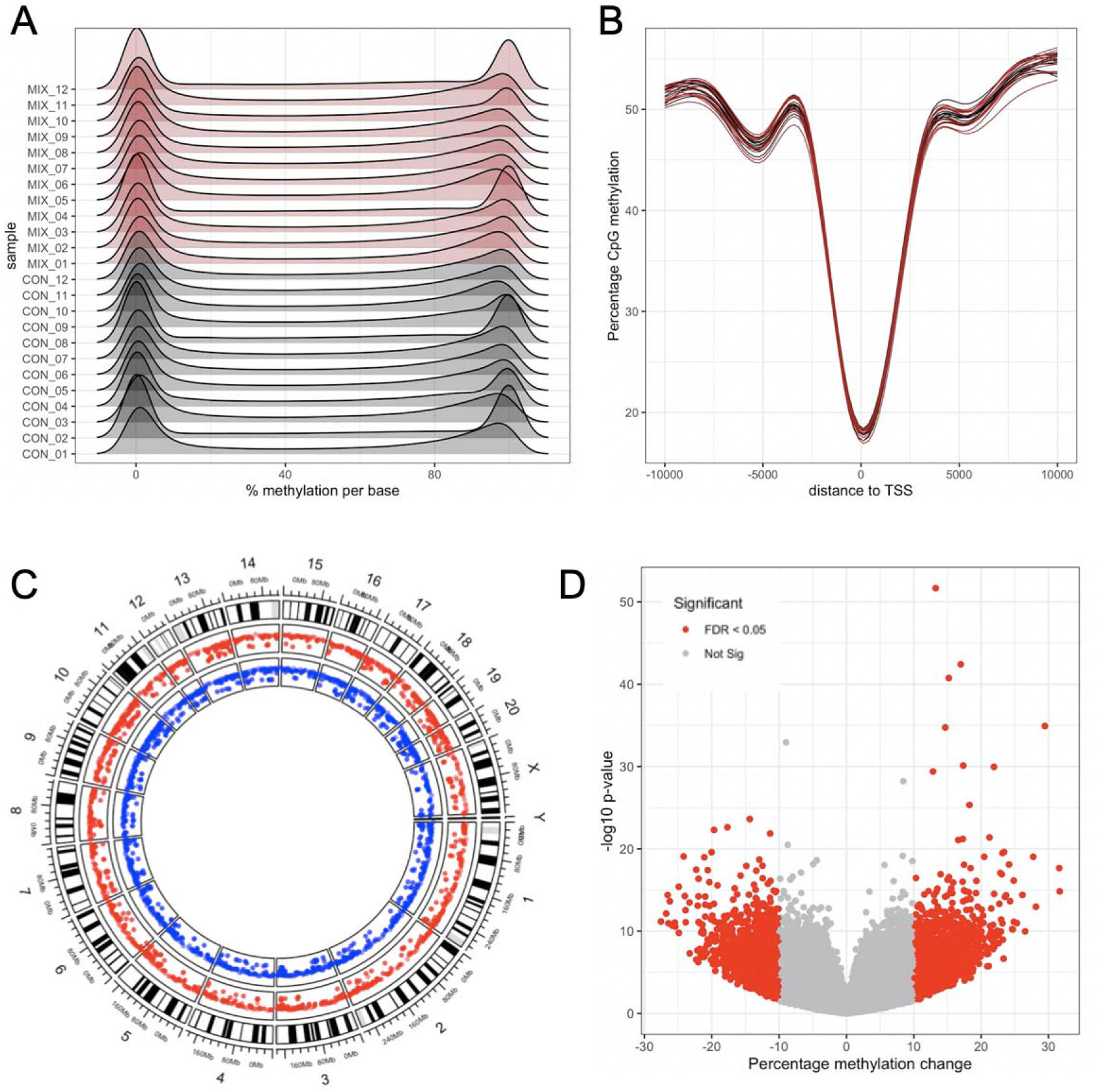
Reduced Representation Bisulfite Sequencing of liver samples from Sprague-Dawley rats exposed for 90 days to the mixture of 6 pesticides. **A.** Percentage DNA methylation profile shows a bimodal distribution of methylation calls for each sample. **B.** CpG methylation decreases around transcription start sites. **C.** Circos plot shows that differentially methylated CpG sites (blue track, hypomethylated; red track, hypermethylated) are scattered around the genome. **D**. Volcano plots of differentially methylated CpG sites shows that a large number of CpG loci are differentially methylated across the rat liver genome with moderate methylation changes.

RRBS of the liver samples was also performed to assess if alterations in epigenetic (DNA methylation) status may be responsible at least in part for the treatment-related changes in gene expression patterns. There were no differences in the percentage of methylated cytosines in CpG islands (42.0 ± 1.6% for controls vs 42.2 ± 2.2% for pesticide-exposed rats). We also analysed the distribution of the percentage methylation per base. Since a given base is generally either methylated or not in a given cell, it is expected that the distribution of percentage DNA methylation per base has a bimodal distribution, which is the case in our study (Figure 4A). We also annotated the methylation calls. DNA methylation patterns in relation to gene transcription start sites (TSS) showed that CpGs near TSS tend to be unmethylated, which confirmed that our RRBS analysis is of a good quality (Figure 4B). We identified 4255 differentially methylated CpG sites (FDR < 0.05) with a modest methylation difference (> 10%) between the group of rats exposed to the pesticide mixture and the control animals (Figure 4C and 4D, (Excel Table S7)). They were mostly located in intergenic regions (50.1%) and introns (37.0%), and to a lesser extent in exons (6.2%) and promoters (6.6%), at an average distance of 56kb from TSS. Only 114 CpG sites presented differences in methylation levels over 25% (max 31.6%). A total of 24 differentially methylated CpG sites were present in promoters (Table 3). Interestingly, the lowest p-value was for hypermethylation of the promoter of the gene coding Coenzyme Q10A, a protein attenuating high-fat diet-induced non-alcoholic fatty liver disease (Chen et al. 2019). We also evaluated the correlation between gene expression and DNA methylation. There was no correlation between the fold changes in gene expression and percentage methylation changes and it was thus not possible to attribute changes in gene expression caused by the mixture to change in DNA methylation at their promoters (Figure S3). However, as gene regulatory elements can be present within introns and as distantly located enhancers, changes in DNA methylation status in non-promoter regions as we have found can also in principle impact their function and alter levels of expression.

Overall, exposure to the mixture of pesticides tested altered the gut-liver axis and caused changes in metabolites from the tryptophan-nicotinamide conversion pathway, which reflects a metabolic adaptation that may exceed cellular capacity for homeostasis, ultimately leading to disease.

## Discussion

We report here the first direct comparison of standard histopathology and serum biochemistry measures and multi-omics analysis of multiple physiological compartments of rats exposed to a mixture of six pesticides (azoxystrobin, boscalid, chlorpyrifos, glyphosate, imidacloprid, thiabendazole) that are most frequently found in EU foodstuffs. Each pesticide was administered at its regulatory permitted EU ADI and thus the expectation was that no effects or signs of toxicity would be observed. However, our results show that the low dose mixture of pesticides we tested caused metabolic alterations in the caecum and blood metabolome, with consequences on liver function, which was mostly reflected by changes in the conversion of tryptophan to nicotinamide. Notably, these metabolic alterations were not detected by regular clinical and biochemical analyses as currently recommended in OECD guidelines and required by government regulatory agencies, but with the new generation of high-throughput ‘omics’ methodologies.

Unlike standard blood biochemical and organ histological analysis, an in-depth molecular profiling using a combination of high-throughput ‘-omics’ methods revealed metabolic effects of the mixture of six pesticides. Histological analysis showed a nonsignificant increase in liver and kidney lesions (Figure 2A and 2B). Considering the relatively short duration of exposure (90 days), and the limited number of animals per test group, we cannot rule out the possibility that these non-statistically significant increases were attributable to the treatment. Animal bioassays are generally extended to 12 months and performed on a larger number of animals (OECD Test Guideline 452: Chronic Toxicity Studies) to detect chronic toxicity from exposure to a given chemical. Contrastingly, the use of more comprehensive ‘omics’ technologies such as metabolomics allowed us to describe in detail the metabolic effects of the pesticide mixture even after just 90 days of exposure. More studies are needed to determine if a longer treatment period with the pesticide mixture leads to adverse health effects. Nevertheless, our results show that the inclusion of ‘omics’ high-throughput approaches in the battery of tests used to study the effects of chemicals promises to substantially heighten their sensitivity and accuracy. This in turn will enhance the ability of such tests to predict risks of toxicity for regulatory decision-making purposes (Hewitt and Herget 2009; Rovida et al. 2015). This could ultimately reduce the duration of animal bioassays and the number of animals needed, which would be in accord with ongoing efforts to improve animal welfare in research and testing.

Although caecum metabolomics showed that there was an effect on gut microbial pathways involved in tryptophan and serotonin metabolism (Table 2), shotgun metagenomics failed to show an effect on gut microbiome functional potential (Figure 3). Gut microbiome metagenomics and metabolomics are part of an emerging field of research and results from studies can be confounded by a large number of factors (McLaren et al. 2019). This could be amplified in our case by the housing of three rats per cage, since rats are coprophagous and thus exchange their gut microbiota (Suckow et al. 2005). It is also not clear if the gut microbiome is mediating the effects of this pesticide mixture, or if the change in liver metabolism is influencing gut microbiome composition and function. The gut microbiome and the liver evidently influence each other, and a large number of studies have shown a bidirectional communication involving bile acids, antimicrobial molecules or dietary metabolites (Tripathi et al. 2018). It is also possible that the pesticide mixture tested here had an effect on bacterial metabolism, which did not affect growth properties. This possibility is supported by a recent study of the soil filamentous fungus *Aspergillus nidulans* (Mesnage et al. 2020). This study found that exposure to the herbicide Roundup GT+ caused alterations in secondary metabolism at a concentration that, however, caused no change in growth.

Changes in caecum and serum metabolome and liver transcriptome profiles were very consistent and suggested that the mixture of pesticides tested in this study triggered a cell danger response (CDR). The CDR is a metabolic response to a physical or chemical hazard, which exceeds cellular capacity for homeostasis (Naviaux 2014). CDR involves the production of reactive oxygen species (ROS), the development of inflammation, and the activation and engagement of detoxification mechanisms. Low levels of pyridoxal are known to be associated with changes in tryptophan levels, as we observed in our study, in order to sustain an active CDR and cause inflammation (Naviaux 2014; Paul et al. 2013). This also explains the increase in nicotinamide levels, which is considered to be a marker for the generation of ROS (Schmeisser et al. 2013). Nicotinamide is a form of vitamin B3, which can act as a stress signal mediating compound, released when DNA-strand breakage is caused by oxidative damage (Berglund 1994). Nicotinamide has been shown to protect from hepatic steatosis by increasing redox potential (Ganji et al. 2014). An improvement in hepatic transaminase concentration is detectable when hypertriglyceridemic patients are treated with nicotinamide (Hu et al. 2012). Overall, this suggests that the increase in nicotinamide levels we observe reflects a metabolic adaptation to oxidative stress induced by exposure to the mixture of pesticides.

Given the known roles of pyridoxal (a form of vitamin B6) and serotonin in autism-related social behaviors (Adams et al. 2006; Muller et al. 2016), the effects of the pesticide mixture on altering the levels of these compounds in our study could point to a role of pesticide exposure in the development of autism spectrum disorder. Whereas we observed an increase in both pyridoxal and serotonin in the caecum we found a decrease in the level of pyridoxal in serum suggesting that the pesticide mixture could lead to a deficiency in vitamin B6. In a recent study, administration of a high-dose of glyphosate caused autism-like behaviors in murine male offspring (Pu et al. 2020). Chlorpyrifos has also been found to be associated with an increased risk for autism spectrum disorder with intellectual disability in human populations (von Ehrenstein et al. 2019). Our observations suggest that further studies are warranted to evaluate the effects of the pesticide mixture we have tested with exposure initiated prenatally in laboratory animals to assess its contribution to the development of behavioural disorders.

The changes in serum metabolite levels we see in this study do not reflect a diseased state, but probably rather metabolic adaptation, which can lead to the development of a pathological state if the damage produced exceeds the capacity for repair. The conversion of tryptophan to nicotinamide is known to decrease in rats presenting a steatotic liver (Ganji et al. 2014), which is the opposite to what we observed. Similarly, pipecolate levels are known to be increased in patients with liver disease, with the level of increase being proportional to the severity of liver damage (Kawasaki et al. 1988). In our study, pipecolate serum levels displayed the opposite trend; that is, they were decreased in animals exposed to the pesticide mixture (Table 2). This could be related to hormesis, a phenomenon by which mild induced stress can give rise to a positive physiological counter-response inducing maintenance and repair systems (Rattan 2013). This is well described for the effect of pesticides for both target (Tang et al. 2019) and non-target (Calabrese and Baldwin 2003) species. Other known hormetic stressors include exercise (Nikolaidis et al. 2008) and fasting (Mattson 2012; Mesnage et al. 2019). Although physiology may initially improve by initiating protective measures in the face of exposure to mild stressors, this can ultimately give rise to a pathological status if the intensity of the stimulation exceeds cellular capacity for homeostasis (Andersen et al. 2005). Although rats exposed to the pesticide mixture in this study presented more signs of pathology than animals in the control group, a longer period of exposure would be needed to determine if liver and/or kidney function will be impaired.

Human populations are exposed to a large range of chemicals on a daily basis. Thus, chemical toxicity tests administering a single substance at medium to high doses in laboratory animals does not reflect a real-life exposure scenario. Other studies have demonstrated toxic effects from mixtures of chemicals at concentrations below regulatory levels (Docea et al. 2018; Lukowicz et al. 2018). It is not clear if the effects detected in our study are due to an interaction between all six components of the pesticide mixture, or if they result from either a single compound or sub-set of compounds. In addition, future studies could explore a dose-response of the effects we observed in both sexes, which would be important in establishing a ‘no observed adverse effect level’ of exposure on the basis of which a safe ADI can be calculated for this pesticide mixture.

In conclusion, our study reveals that metabolic effects following exposure to a pesticide mixture, which can be typically found in EU foodstuffs, and administered at the ADI, can be detected using caecum and serum metabolomics, and liver transcriptomics and DNA methylation profiling. Crucially, these molecular biological and metabolic changes were not detectable using conventional biochemical and histopathological investigations, which regulators currently rely upon for chemical risk assessment. Thus our results highlight the advantages of incorporating high throughput ‘-omics’ methods into OECD Guidelines for the Testing of Chemicals (Sauer et al. 2017). Although additional studies are needed to determine if longer exposure to the pesticide mixture we tested leads to adverse effects, our results demonstrate that high-throughput ‘omics’ analyses as applied herein can reveal molecular biological and metabolic biomarkers, which can potentially act as more sensitive and accurate predictors of long-term health risks arising from pesticide exposures. This in turn can lead to more appropriate regulatory public health protection measures.

## Acknowledgements

This work was funded by the Sustainable Food Alliance (USA), and in part by the Franciscan Health Foundation (USA) and the Sheepdrove Trust (UK), whose support is gratefully acknowledged.

## Competing interests

RM has served as a consultant on glyphosate risk assessment issues as part of litigation in the US over glyphosate health effects. The other authors declare no competing interests.

