## Supplementary Material for "Multi-omics phenotyping of the gut-liver axis allows health risk predictability from *in vivo* subchronic toxicity tests of a low-dose pesticide mixture"

Robin Mesnage, et al.,

**Table of Contents**

Table S1. Bacterial strains used in this study.

Table S2. Analysis of our transcriptome findings with the drugMatrix toxicogenomics database.

Figure S1. Effects of all the actives principles alone on the *in vitro* bacterial growth of two different species: EC4 and LB5.

Figure S2. Effect of DMSO on bacterial growth of 10 strains of *L. rhamnosus* and 9 strains of *E. coli*.

Figure S3. No correlation between the fold changes in gene expression and percentage methylation changes.

**Additional document –**

Excel File

- Table S1 – Results of the serum metabolomics analysis
- Table S2 – Results of the caecum metabolomics analysis
- Table S3 – Results of the shotgun metagenomics analysis with IGGsearch
- Table S4 – Results of the shotgun metagenomics analysis with Kaiju
- Table S5 – Results of the shotgun metagenomics analysis with Metaphlan2
- Table S6 – Results of the liver transcriptomics analysis
- Table S7 – Results of the RRBS analysis

File S1.pdf – Feed analysis to identify possible contaminants.

**Table S1.** Bacterial strains used in this study.

| Collection ID | Other ID | Code | Species | Origin |
| --- | --- | --- | --- | --- |
| UCMA n°2926 | ATCC 9595 | LB1 | *Lactobacillus rhamnosus* | Human, saliva |
| UCMA n°2927 | CIP A158 | LB2 | *Lactobacillus rhamnosus* | Human |
| UCMA n°2929 | CIP 71.38 | LB3 | *Lactobacillus rhamnosus* | Human, saliva |
| UCMA n°2930 | CIP 103700 | LB4 | *Lactobacillus rhamnosus* | Human, encarditis |
| UCMA n°2933 | CIP 103888 | LB5 | *Lactobacillus rhamnosus* | Human, liver abscess |
| UCMA n°2934 |  | LB6 | *Lactobacillus rhamnosus* GG | Human, faeces |
| UCMA n°5164 |  | LB7 | *Lactobacillus rhamnosus* | Milk |
| UCMA n°2935 | ATCC 7469 | LB8 | *Lactobacillus rhamnosus* | Human |
| UCMA n°20972 | CIP 104456 | LB9 | *Lactobacillus rhamnosus* | Human, faeces |
| UCMA n°20973 | CIP 102102 | LB10 | *Lactobacillus rhamnosus* | Human, faeces |
| UCMA n°6835 | CIP 53.126 | EC1 | *Escherichia coli* | Human, faeces |
| UCMA n°7218 |  | EC2 | *Escherichia coli* O157:H7 | Heifer, faeces |
| UCMA n°7733 | ATCC 10798 | EC3 | *Escherichia coli* K-12 | Human |
| UCMA n°9748 | DSM 6601 | EC4 | *Escherichia coli* nissle 1917 | Human |
| UCMA n°20974 | CIP 105182 | EC5 | *Escherichia coli* O159:H34 | Human, faeces |
| UCMA n°7088 |  | EC6 | *Escherichia coli* | Rotten milk |
| UCMA n°7105 |  | EC7 | *Escherichia coli* | Camembert cheese |
| UCMA n°7217 |  | EC8 | *Escherichia coli* | Milk |
| UCMA n°10529 |  | EC9 | *Escherichia coli* | Raw milk |

**Table S2. Comparison of our transcriptome findings to a list of gene expression signatures collected from various rat tissues after treatments with various drugs from the drugMatrix toxicogenomics database.**

| Term | Overlap | P-value | Adjusted P-value | Genes |
| --- | --- | --- | --- | --- |
| Niacinamide-750 mg/kg in Water-Rat-Liver-3d-up | 22/309 | 1.96E-11 | 1.54E-07 | SRRM2;GDF15;PLK2;TAT;MIF;OPLAH;CLU;AGT;PNRC1;VTN;PSMC5;CYP2C22;CPT2;PSMC4;BAG3;CBS;HSD17B2;PSMC2;CFL1;PMVK;PGK1;JUNB |
| Ipriflavone-1500 mg/kg in CMC-Rat-Liver-3d-up | 20/318 | 1.47E-09 | 5.79E-06 | CDKN1A;GDF15;PLK2;TAT;MIF;NR0B2;AGT;PNRC1;HEBP1;PDLIM1;CYP2C22;CPT2;PSMC4;BAG3;HSD17B2;PSMC2;CFL1;PGK1;MLYCD;JUNB |
| Letrozole-250 mg/kg in Corn Oil-Rat-Liver-5d-up | 20/319 | 1.55E-09 | 4.08E-06 | CDKN1A;TSC22D3;TAT;MIF;CLU;AGT;PNRC1;PDLIM1;RGD1309534;CYP2C22;PSMC4;BAG3;CBS;PSMC2;CFL1;ESD;PGK1;ENPP3;JUNB;UOX |
| Marimastat-1000 uM in DMSO-Rat-Primary rat hepatocytes-0.67d-up | 18/276 | 5.69E-09 | 1.12E-05 | CDKN1A;SLC20A1;GDF15;PLK2;TAT;MRPS18B;LITAF;ZFP36L2;PNRC1;VEGFA;RBM3;PDLIM1;PRPF39;RABEP1;PSMC4;PSMC2;PGK1;JUNB |
| Valproic Acid-1500 mg/kg in Water-Rat-Liver-3d-up | 18/293 | 1.44E-08 | 2.27E-05 | SRRM2;ACY1;GDF15;TAT;MRPS18B;MIF;NR0B2;CLU;ZFP36L2;PDLIM1;CYP2C22;CPT2;PSMC4;HSD17B2;PSMC2;PMVK;ESD;MLYCD |
| Phenylhydrazine-78 mg/kg in Water-Rat-Liver-3d-up | 19/335 | 2.06E-08 | 2.70E-05 | CDKN1A;CEBPD;SLC20A1;GDF15;TAT;MIF;LITAF;AGT;ZFP36L2;PNRC1;VEGFA;PDLIM1;VTN;PSMC4;HSD17B2;PGK1;DNAJB9;JUNB;UOX |
| Sulfaphenazole-1695 mg/kg in Water-Rat-Liver-1d-up | 17/282 | 4.85E-08 | 5.46E-05 | ACY1;TAT;MIF;NR0B2;CLU;AGT;DPP4;PDLIM1;PSMC5;CYP2C22;CPT2;PSMC4;CBS;HSD17B2;PSMC2;PGK1;ENPP3 |
| Mifepristone-3 mg/kg in Corn Oil-Rat-Liver-3d-dn | 18/319 | 5.31E-08 | 5.23E-05 | SRRM2;PLK2;TAT;MIF;NR0B2;CLU;PNRC1;VEGFA;PDLIM1;VTN;CYP2C22;CPT2;PSMC4;CBS;HSD17B2;PSMC2;ESD;PGK1 |
| Olanzapine-23 mg/kg in CMC-Rat-Liver-3d-up | 18/335 | 1.11E-07 | 9.71E-05 | ACY1;GDF15;PLK2;TAT;MIF;NR0B2;AGT;PNRC1;HEBP1;VTN;RGD1309534;CYP2C22;PSMC4;BAG3;PSMC2;ENPP3;JUNB;UOX |
| Aminoglutethimide-350 mg/kg in CMC-Rat-Liver-1d-up | 17/307 | 1.65E-07 | 1.30E-04 | ACY1;TAT;MRPS18B;MIF;CLU;PDLIM1;PSMC5;RGD1309534;CYP2C22;CPT2;PSMC4;CBS;HSD17B2;PSMC2;CFL1;ESD;PGK1 |

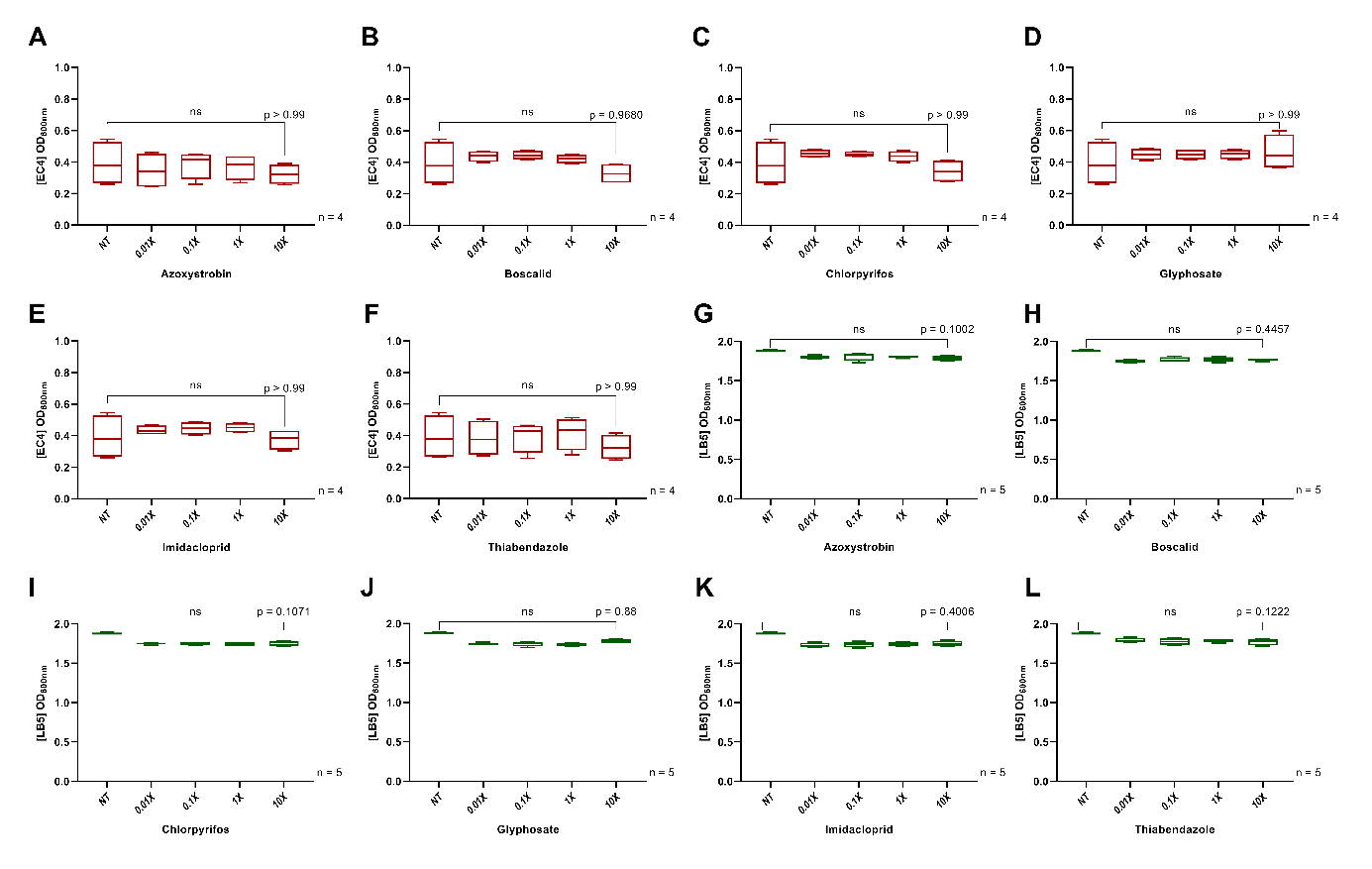
 **Figure S1. Effects of all the actives principles alone on the *in vitro* bacterial growth of two different species: EC4 and LB5.** The bacterial growth of the species *E. coli* (EC4) **(A-F)** is not impacted by any active principles of pesticides tested azoxystrobin **(A)**, boscalid **(B)**, chlorpyrifos **(C)**, glyphosate **(D)**, imidacloprid **(E)** and thiabendazole **(F)**. In the same way, the bacterial growth of the species *L. rhamnosus* (LB5) is not significative impacted by one of the active principles **(G-L)**.

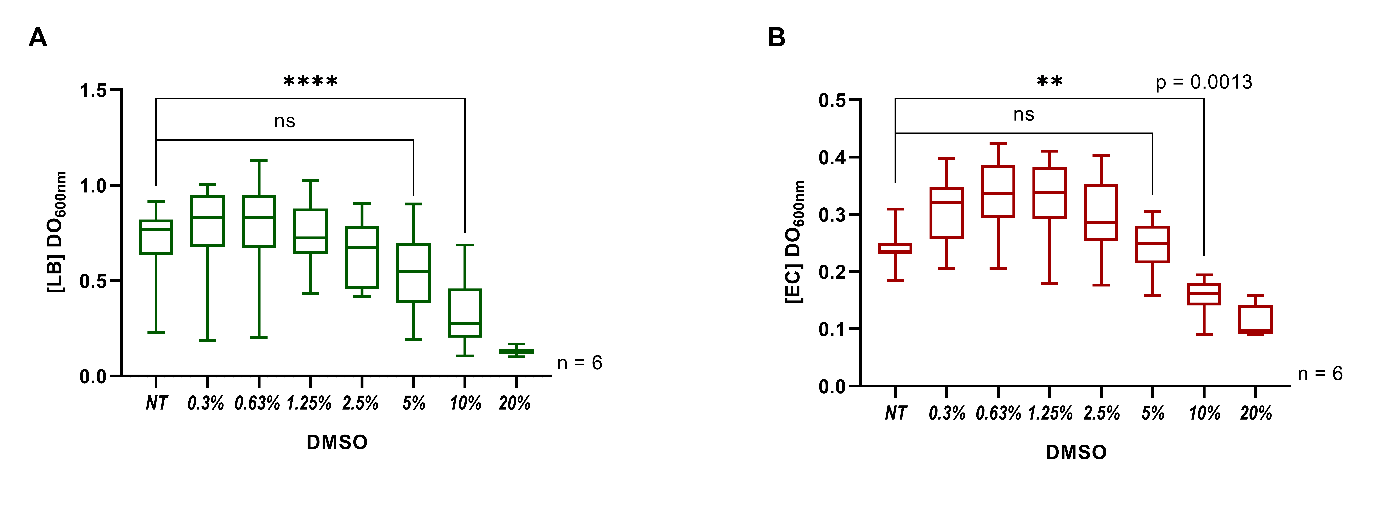

**Figure S2. Effect of DMSO on bacterial growth of 10 strains of *L. rhamnosus* and 9 strains of *E. coli*.** The bacterial growth of 10 strains of *L. rhamnosus* **(A)** and 9 strains of *E. coli* **(B)** is not significantly impacted under a concentration of 5% DMSO. Under 1% DMSO, all bacterial strains have growth equivalent to the control (NT).

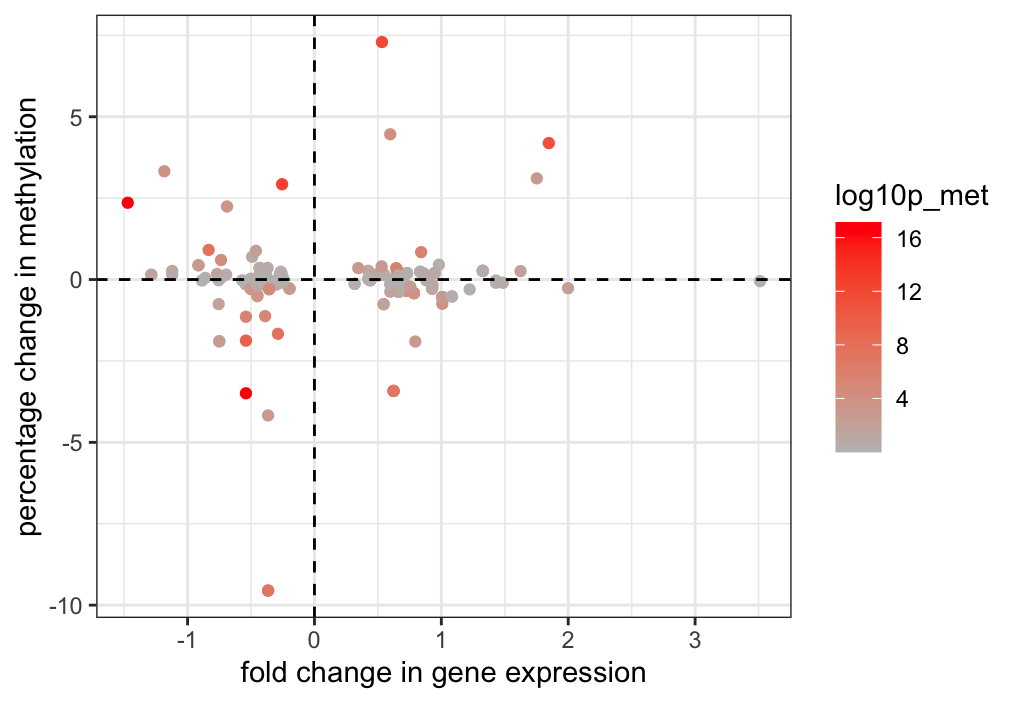

**Figure S3. No correlation between the fold changes in gene expression and percentage methylation changes.** RRBS of the liver samples was performed to assess if alterations in epigenetic (DNA methylation) status may be responsible at least in part for the treatment-related changes in gene expression patterns. The fold changes in gene expression (RNA-seq) are compared to the percentage of change in methylation (RRBS) for the differentially expressed genes.
